# In the Line of Fire: Debris Throwing by Wild Octopuses

**DOI:** 10.1101/2021.08.18.456805

**Authors:** Peter Godfrey-Smith, David Scheel, Stephanie Chancellor, Stefan Linquist, Matthew Lawrence

## Abstract

Wild octopuses at an Australian site frequently propel shells, silt, and algae through the water by releasing these materials from their arms while creating a forceful jet from the siphon held under the arm web. These “throws” occur in several contexts, including interactions with conspecifics, and material thrown in conspecific contexts frequently hits other octopuses. Some throws appear to be targeted on other individuals and play a social role, as suggested by several kinds of evidence. Such throws were significantly more vigorous and more often used silt, rather than shells or algae, and high vigor throws were significantly more often accompanied by uniform or dark body patterns. Some throws were directed differently from beneath the arms and such throws were significantly more likely to hit other octopuses. Throws targeted at other individuals in the same population, as these appear to be, are the least common form of nonhuman throwing.

## INTRODUCTION

The throwing of objects is an uncommon behavior in animals. Throwing has sometimes been seen as distinctively human, and it probably does have an important role in hominin evolution (1, 2). But targeted throwing has also been observed in some non-human primates (especially chimps and capuchins), elephants, mongooses, and birds (3–5). Many, though not all, of these observations were made in captivity, or with populations acclimated to human presence. Other behaviors related to throwing include the flicking of irritating hairs at threats by spiders, the squirting of water at prey by archerfish, and the swinging of silk threads at prey by some bolas spiders (6–8). Antlion larvae fling sand when prey is attempting to escape from their conical traps. Throws are oriented towards the direction of the prey (9), though the throws may function as much to destabilize the walls of the trap as to disrupt behavior of the prey itself. The processing of prey by various predators, such as dolphins, also features thrashing and tossing (10).

Directed throwing can target prey or encased food (seen in birds and mongooses), threats or problematic animals of other species (elephants, various primates) and more rarely, other individuals of the same population (chimps) (1–5). Throwing of this last kind is simultaneously a form of tool use and a social interaction; a projectile is an agonistic social tool. In all non-human cases, it is difficult to show deliberate targeting. The basis for a claim of targeting has often been the fact of regular hits on ecologically significant objects, along with apparent attention and aiming, although the use of these criteria is often implicit.

Octopuses are dextrous manipulators of objects. Veined octopuses (*Amphioctopus marginatus*) carry shelter in the form of nested half-shells that are then reassembled (11). Octopus den-building and maintenance includes the use of diverse materials, as seen in the use of a collected piece of dead sponge to block a den entrance (12). Octopuses have often been seen as largely asocial animals, but recent work has shown tolerance, signal use and other socially-directed behaviors in some species (13, 14).

Here, we describe a complex behavior by wild octopuses involving coordinated use of the arms, the web, and jets of water from the siphon, that results in material being forcibly projected through the water column, sometimes hitting other octopuses. We compare this behavior – a jet-propelled throw – to other octopus behaviors, examine the roles it has in social and non-social contexts, and consider whether some throws are targeted on other individuals.

## RESULTS

The observations described here were collected at a field site where unusually high numbers of *Octopus tetricus* interact within a small area of approximately 8 m^2^, at the time of data collection (15, 13). The data used here comprise one full day and two adjacent part days of continuous daylight filming in January 2015, along with a shorter continuous period of filming from a single day in December 2016, marked by notable examples of the patterns we describe using the 2015 data. The 2015 data comprises 20 hours 37 mins of video in total, of which 13 hours 29 minutes were recorded on the middle full day. In that 2015 data, from four to as many as eight individuals were visible at a time at the site and engaged in 1316 interactions. The eight octopuses observed as a snapshot maximum were probably (see Methods) three males and five females. During many periods, there was a particularly active male present (the “most active male,” below), plus several females, and often an additional peripheral male. The most active male interfered with approaching and departing octopuses to various degrees.

The average hourly high count (the maximum for each hour, averaged) on the day with the largest sampling was 6.3 individuals (s.d.=0.89). Interactions were frequent; from 11 up to 234 per hour, average of 73 (s.d. 60.96; see SI). These included fights of various levels of intensity, matings, and also approaches or reaches from one octopus to another, who then reacted (with color or posture change, another reach, ducking, retreat, etc).

A conspicuous behavior observed in the video data involved the coordinated use of arms, web, and siphon, by which material was projected through the water column (Figure 1 A and B). In this behavior, octopuses gather material in their arms, hold it in the arm web, and then use the siphon to expel the material under pressure. We could seldom view all mechanics of these actions on a video, but typical cases proceed as follows: after the gathering of material, commonly from inside the den, the siphon is brought under the web of the octopus’ arms, by bringing it between arms L4 and R4 (the two rear-most arms), and water is expelled forcibly as the material held is released (see Figure 1 C and D). The material is ejected from between arms L1 and R1 (the two front-most arms) in the majority of throws, but in some cases the material was ejected between other adjacent pairs of arms, or directly under one arm. These jets may be strong enough to propel material several body-lengths from the animal in still water, or so weak that the material falls almost directly in front of the animal. In some cases the projected material hits another octopus, or another possible target.

**Figure 1:**
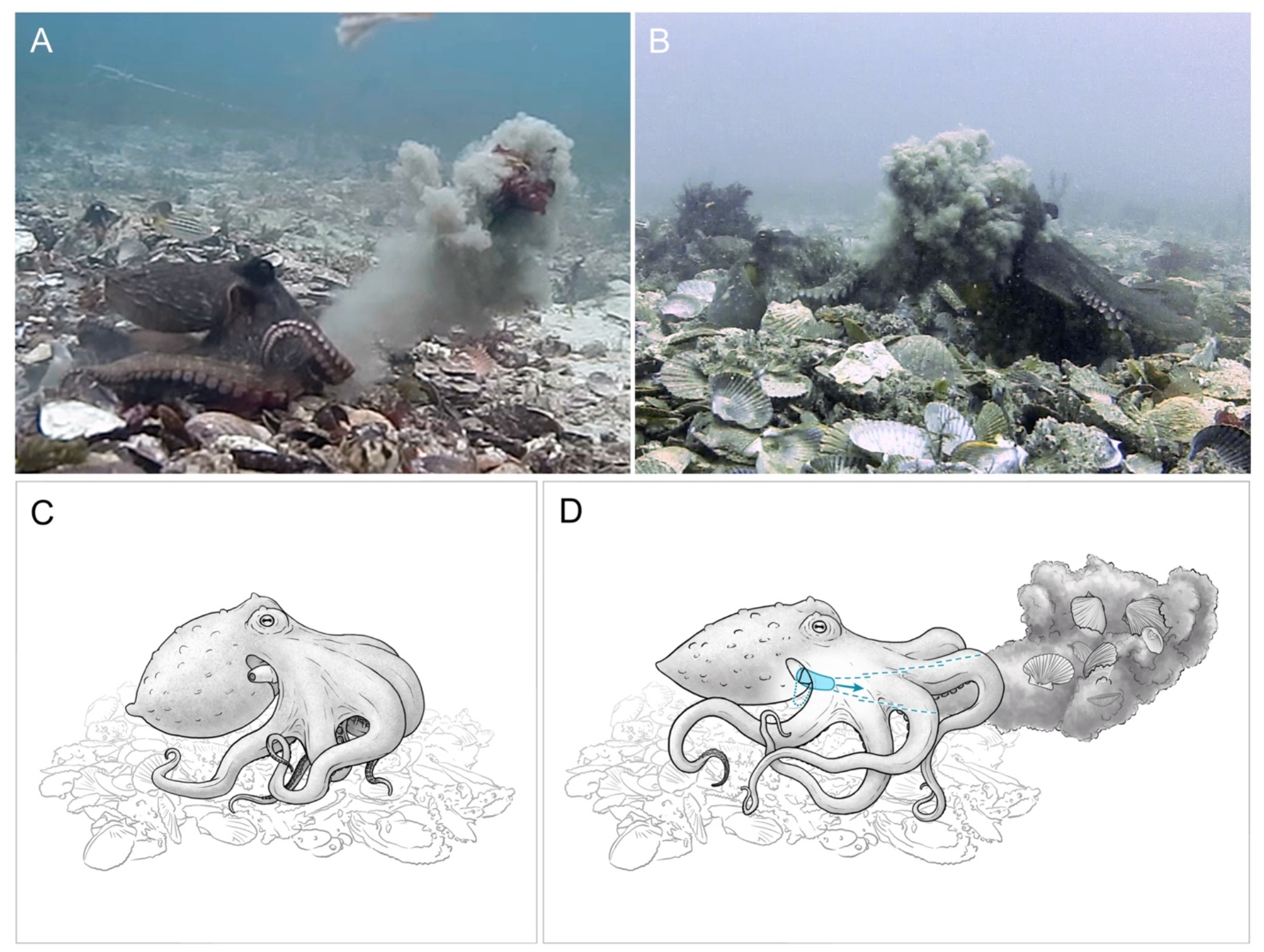
Debris throwing by *Octopus tetricus* in the wild: Panel A - Octopus (left) projects silt and kelp through the water (from video by Peter Godfrey-Smith); B – an octopus (right) is hit by a cloud of silt projected through the water by a throwing octopus (left; see SI for video of this event); C, D The mechanics of throwing behavior, C – shells, silt, algae or some mixture is held in the arms preparatory to the throw, mantle is inflated preparatory to ventilation during the throw, siphon at this stage may still be visible in its usual position projecting from the gill slit above the arm crown; D – siphon is brought down over rear arm and under the web and arm crown between the rear arm pair (arms R4 and L4), and water is forcibly expelled through the siphon, with contraction of the mantle, as held debris is released, projecting debris through the water column. Illustrations by Rebecca Gelernter.

We interpret these behaviors as *throws*. This behavior contrasts with pushing material using the arms. The previously described “Remove” behavior (16) broadly construed – using the jet on material regardless of whether it is gathered and held – includes throws in our sense, along with non-throwing behaviors in which a water jet is directed on an object. A clear case of a throw in our sense requires a water blast from the siphon that is directed at and simultaneous with release of material held in the arms, requiring that the siphon move into an unusual position below the arm web.

Throwing by octopuses is a common behavior at this site and also occurs at another site (17). At this site, throwing occurred in 11 of 12 sampling periods which spanned eight years (2011 to 2018). In this report, we closely examine throws in sampling periods during 2015 and 2016.

Throws in this sense do shade into other behaviors, as the gathering and holding of material can be minimal. Octopuses have several other ways of moving debris, including pushing, scooping from below, and picking-and-dropping. We include in our dataset here borderline cases where a jet was used from below but the gathering of material was minimal (these cases comprise about 11% of the total; see SI) along with one unusual case where a shell was, at least in part, flung by straightening an arm, and hit another octopus. We do not count cases where material propelled is clearly not held, and/or the siphon is directed on material while the siphon is above the arm web. For brevity, we refer to all items in the data set as “throws,” where this should be understood to include borderline cases.

### Throws by different individuals and sexes

We observed N=102 of these throw behaviors in the 2015 video data. A few individuals accounted for most of the throws, but likely half or more of individuals present threw at least occasionally. Tracking of individuals is difficult at this site, as octopuses frequently enter and leave the area under video coverage. They also change dens. A few octopuses had identifying marks repeatedly visible from video. The maximum number present at a time, as noted above, was 8. We estimate, based on diver counts of numbers both on and around the site during the period of data collection, that the total number of individual octopuses present in the 2015 data was around 10.

Out of 102 throws in the 2015 data, two reliably reidentified females were responsible for 67 throws (41 by octopus T23F and 26 by T1F). For both T23F singly and both together, this was significantly more than expected under a binomial distribution, assuming four or more potential throwers (p=<0.001). These two individuals were responsible for at least 66% of all throws, and likely another 5 throws when identifying marks were not visible on video to confirm a suspected ID. The most active male mentioned above may have been the next most frequent thrower, with 5 throws in total, though he could often only be provisionally reidentified until he was bitten by a fish midway through the recording period, leaving a visible scar.

The distribution of N=29 throws among other individuals is uncertain; using strict identification criteria, there were 24 *nominal* individuals (including the three described above) who engaged in throws across the three days sampled in 2015, where a nominal individual is an octopus continuously on-screen or with unique markings (7 females identified by mating behaviors, 7 by other behaviors (see SI); 7 males identified by mating, 2 by other behaviors; one with no assigned sex). The actual number of individuals who spent time on the site was, as noted, probably much smaller and around ten, with eight the largest number present at one time; the count of nominal individuals assumes in effect that every break in onscreen continuity in the absence of distinguishing marks establishes a new individual.

There were 90 throws by females and eleven by males, a ratio of 8.9:1. The female to male sex ratio at the site probably ranged from around 1.7:1 to around 5:1. This ratio varied continually as octopuses came and went and could not be precisely determined at each moment from the video. It was affected by apparent attempts by the most active male to alter the movements of other octopuses.

A large number of throws were also made by octopuses while at a particular den. The most frequent throwers T1F and T23F each occupied this den at different times, as did other individuals briefly, and in total 59 throws (58%) originated at this location (57 throws by the two most frequent throwers). This den was closely flanked by other octopus dens, one at 20 cm, was close to a camera, and was frequently visited by other passing octopuses.

On the day with 13.5 hours of sampling, throws occurred throughout daylight hours but increased in frequency around dusk (see SI). This timing coincided with the largest number of interactions in an hour, as well as the largest count of octopuses simultaneously in frame, At dusk, octopuses remained active until the video became too dark to evaluate.

### Context of behaviors, and materials thrown

We distinguished three contexts in which throws occurred. *Eating* throws discarded the remains of a meal after foraging and eating (the duration of which was variable but could exceed 30 minutes). *Den maintenance* throws were preceded by rearranging or excavation of materials from in or around the den. We categorized as *social* any throw that occurred during or within a 2 minute window of an interaction with another octopus. As noted above, interactions included fights, mating attempts, and approaches or reaches followed by an apparent reaction by another octopus. Contexts were not mutually exclusive, as, for example, an octopus could interact with another octopus while cleaning its den; we scored these throws as mixed. Other cases were scored as “no context,” when none of these indicators were present. In a small number of cases (N=4) that were difficult to score, events outside the 2 minute window were taken into consideration or events within it set aside (see SI).

N=101 throws were scored for their context in this way (with one throw at the start of sampling not scored because the two minutes preceding the throw were not recorded.) Over half of all throws (53%) occurred in a context that was at least partly social (36 throws in social contexts (36%); 17 in mixed contexts combining social and den-cleaning (17%); 32 during den-cleaning (32%); 8 after eating (8%); and 8 with no apparent context (8%)).

Nearly half of the nominal individuals (11 of 24) threw in social contexts, and thus social throwing was common among octopuses as well as among throws.

Octopuses threw three different kinds of material: shells, silt, and algae. Shells and silt were ubiquitous at all times at the site; algae was not ubiquitous but frequently within reach. In general, we scored a throw according to its main material and discounted minor contributions - for example, small amounts of silt in shell throws. Some throws were scored as mixed (50% per material), however, as there were substantial amounts of two materials. Shells were thrown most often and algae least: N=55 shell throws, 35 silt throws, and 11 algae throws.

The materials thrown differed across social and non-social contexts. In social contexts, octopuses threw silt significantly more often than in other contexts, whereas they threw shells more often in den cleaning contexts (Figure 2 A and B. Chi-square test: N = 76, χ^2^ = 9.26, df = 2, p < 0.01. Only social, mixed and den cleaning contexts were considered due to sample size). Shells were also thrown in all eating contexts, along with algae in one case. The frequency of throwing algae differed little across contexts (Figure 2a).

**Figure 2:**
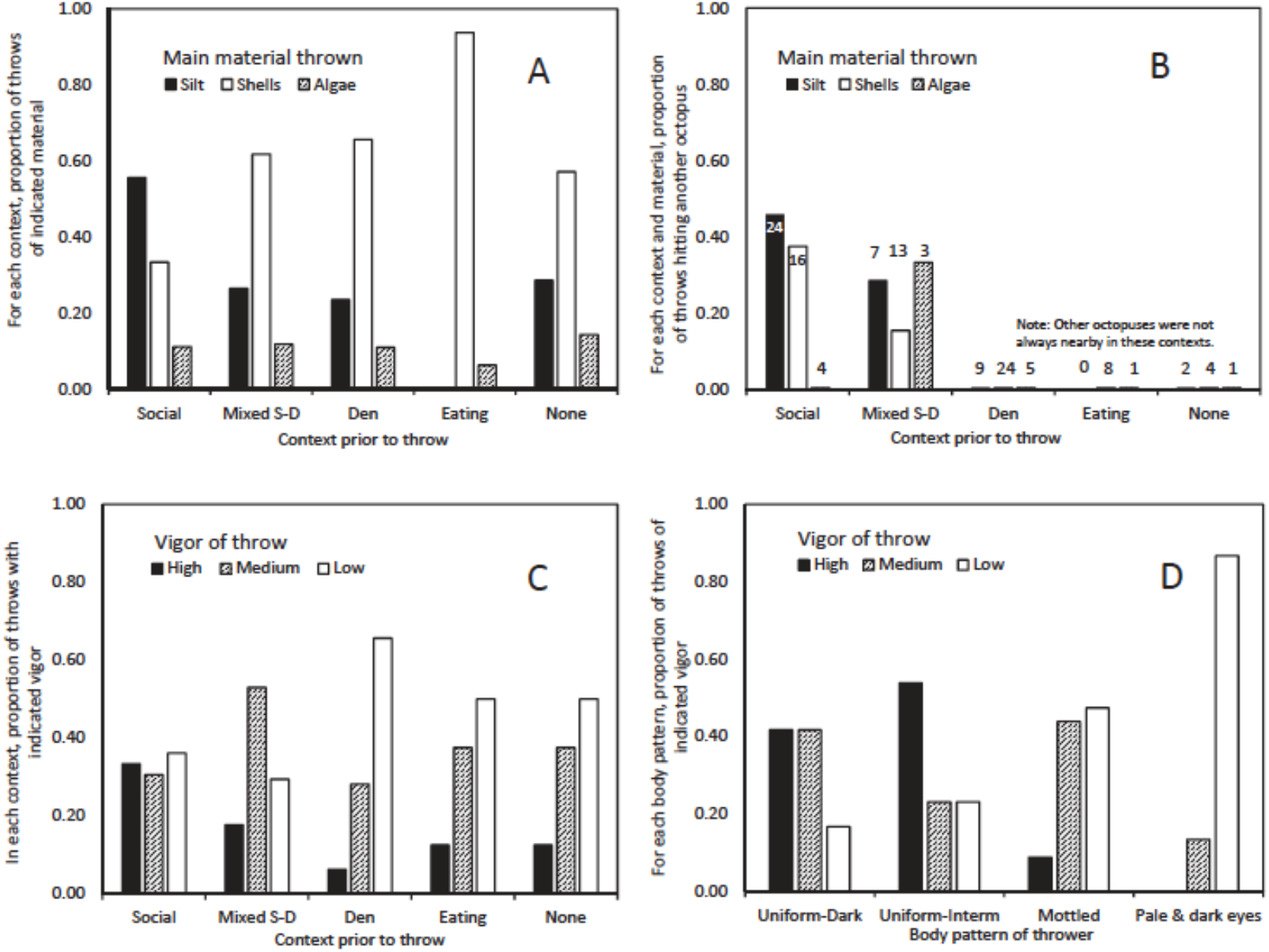
Throwing behavior by *Octopus tetricus* in the wild: Panel A, B of different materials by context (“mixed S-D” indicates both social and den maintenance contexts occurred prior to the throw, see text for details), and C, D thrown with varying vigor. Panel A – the proportion of all throws; B – the proportion of throws that hit another octopus (numbers above columns indicate sample size of throws of that context and material); C – the proportion of throws by context; and D – by body pattern displayed during the throw by the thrower.

The vigor of throws also differed across contexts. Throws were scored as low, medium, or high vigor (see SI for criteria). Overall, low vigor throws were most common. (Low vigor: 48 of 102 throws (47%); medium, 35 throws (34%); high, 19 throws (19%)). However, in social contexts high vigor throws were 12 of 36 total, whereas in den cleaning, high vigor accounted for only 2 of 32, see figure 2c) of throws. Considering just throws in social, mixed or den contexts (those with the greatest sample size), significantly more social throws were high vigor and den cleaning throws were low vigor (Figure 2c. Chi-square test: N=85, χ ^2^= 12.98, df = 4, p = 0.011).

Vigor was also associated with the color of the thrower, at the moment of throw (see figure 2d). Earlier work at this site has found that darker colors are associated with more aggressive behaviors (13). This pattern extends to the vigor of a throw. Octopuses that displayed uniform color (dark or medium) threw significantly more often with high vigor, while those displaying a “pale and dark eyes” pattern threw more often with low vigor, (chi-square test: N=102, χ^2^ = 29.7, df = 4, p < 0.0001). This is probably a form of disruptive coloration (18, 19) though usually only the top of the body could be seen. Those displaying a mottled pattern also threw less often with high vigor.

### Hits on other octopuses

In 17 cases, 33% of the throws in social and mixed-social contexts, the material thrown hit another octopus. In two other cases, a throw hit a fish. Throws that hit another octopus (see SI) tended to differ in several ways from throws that did not. These differences collectively suggest a degree of targeting. First, in a majority of all throws, the material was emitted from between the most frontal arms: L1 and R1. However, throws occasionally emitted material from between arms L1/L2, R1/R2, or directly beneath L1 or R1. A total of 14 throws out of 98 that could be assessed were “anomalous arm” throws of this kind. Of these 14 throws, 6 hit another octopus (43%), as compared to 11 hits from 84 throws (13%) of the L1/R1 type. Anomalous arm throws were then more likely to hit other octopuses than L1/R1 throws (p=0.0145, Fisher’s exact test, 2-tailed). In three cases of hits, a thrower also altered their body orientation towards the apparent target, but these movements were very slight, and the effects of arm choice other than L1/R1 were more marked.

Second, throws by octopuses displaying Uniform body patterns (especially uniform dark patterns) hit other octopuses significantly more often than in other body patterns (p=0.0021, Fisher’s exact test, 2-tailed).

Third, we noted a tendency above for throwers in social contexts to throw silt rather than other materials. This tendency was also seen in the case of throws that hit another octopus. Throws solely or primarily of silt, within social or mixed-social contexts, hit another octopus 8 out of 19 times (42%). In contrast, throws solely or primarily of shells in these contexts hit another octopus 4 out of 16 times (25%). This difference was not statistically significant. (See also Figure 2b for a fuller representation, including throws with mixed materials.) In only one case did a throw of algae (along with shells) hit another octopus. In addition, high vigor throws more frequently hit an apparent target (7 of 19 high vigor throws were hits (37%); 6 of 34 medium vigor (18%) and 4 of 48 low vigor throws (8%)) (p=0.017, Fisher’s exact test comparing high to the combined categories of medium and low vigor throws). High vigor throws have longer and often wider range, so this need not reflect deliberate targeting.

### Sequences of throws with large numbers of hits

A particular sequence of behaviors at the end of the fully sampled day also has circumstantial evidence of deliberate targeting and a social role for throws, due to the number of hits, their circumstances, and accompanying behaviors. In this period of approximately 60 minutes, a single female (T23F) threw 17 times, of which 9 were hits on other octopuses. Of those 9 throws, 8 were hits on a nearby likely female and 1 was a hit on the most active male.

Several of these throws occurred in a context of intermittent reaches and grappling of arms between the thrower and the probable female in an adjacent den. In some cases, the target octopus raised an arm up between itself and the thrower, just before the throw, perhaps in recognition of the act in preparation. These preparatory arm raises were more marked in the later throws of that sequence. Two of the hits within this sequence were throws where material was emitted between arms L1/L2 rather than L1/R1. In these cases, had the material been projected from the L1/R1 position, it would apparently have missed the other octopus, as this choice affected the angle of the throw.

This period in the 2015 data can be compared to another period of frequent throws between two octopuses with frequent hits, in December 2016. In this period of approximately 3 hours 40 mins, during which individuals can be reidentified based on onscreen continuity and marks, a single female threw material 10 times, with 5 of these hitting a male in an adjacent den, who attempted several times to mate with her. The male also threw material either once or twice during that period, once disposing of shells remains after eating (with the other case perhaps similar but unclear). All ten of the female’s throws were entirely or partly silt throws. In one hit, the female’s preparatory motions included a turn towards the male, and material was emitted between arms R1/R2, bringing the male directly into the path of the throw. This sequence is also notable for the behaviors of the male who was the apparent target of the hits. In 4 cases the male ducked during the process of the throw itself, and over the course of the sequence, these movements occurred earlier in relation to the throw. In the first two cases, the target ducked after release of the throw; in the latter two, he ducked before the release, during preparatory motions by the thrower.

One other category of throws in the 2015 data shows some evidence of targeting. In 12 throws, material was thrown in the direction of a nearby stationary camera. Throws often had insufficient force to make the camera an apparent target. In three cases, however, the thrower advanced toward the camera before making the throw, or advanced obliquely and threw from one side (between arms R1 and R2) toward the camera. In two cases (one with the throw between R1/R2), the throw hit the camera tripod. Both of these hits occurred during a data-collection period of roughly 100 minutes when a camera was accidentally placed closer to octopus dens than usual. The usual camera distance is about a meter or slightly more, and this placement was closer by approximately a third of that distance. During this period, the octopus in the den closest to the camera threw material 6 times in the direction of the camera, out of 18 throws during that period, including the two hits.

## DISCUSSION

The throwing of material by wild octopuses is common, at least at the site described here. These throws are achieved by gathering material and holding it in the arms, then expelling it under pressure. Force is not imparted by the arms, as in a human throw, but the arms organize the projection of material by the jet. Earlier descriptions and videos have noted other octopus species using their jets to clear debris from a den area, but directing the jet under the arm-web in the way described here is unusual, especially given the octopus body layout. This behavior has not been noted in ethograms (eg., 16) as distinct from using the jet on loose materials. However, similar ‘throw’ behaviors related to den maintenance has been seen in some other species (DS, personal observation). While observations used in this paper were made at a site where octopuses coexist in close proximity, throws functioning in den den-maintenance and removal of food waste may be expected to occur also in contexts with little or no social interaction.

At the site described here, some throws appear to have a social role. Other octopuses were frequently hit by thrown material. Throws within social contexts tended to be more vigorous, and more often used silt as the thrown material. Within social and mixed-social contexts, silt throws resulted in hits more often than throws of other materials, though this last result was not statistically significant. Hits in many cases occurred within sequences of interactions that featured ongoing mild aggression (arm probes and momentary grappling). Throwing in general is more often seen by females, and we have seen only one hit (a marginal one) from a throw by a male. Octopuses who were hit included other females in nearby dens, and males who have been attempting mating with a female thrower. In one period of interaction, a throw that was a hit was followed by a couple of events where the thrower pushed shells into the den of the same neighboring octopus, without throwing them. In some cases, a thrower released material between or below arms other than the usual forward-facing combination of arms L1 and R1, in a way that increased the chance of a hit. Some throws also included a slight turn by the thrower that also made a hit more likely than it would have been without the turn. Targets sometimes appeared to anticipate a throw in preparation, and ducked in a way that partially or entirely eliminated the impact of the hit. On other occasions, they raised arms in the direction of the thrower but did not duck. All this is evidence that throws in some cases are targeted on other octopuses, and function in the management of social interactions, including sexual interactions.

The case for a social role for throwing is tempered by the fact that there are some things we have *not* seen. We have not seen an octopus who was hit by a throw “return fire” and throw back. We have not seen a hit directly initiating a fight, due to immediate retaliation by the target. The general effects of throws were difficult to assess, and we were hampered in this respect by sample size and limitations on the identification of target octopuses. Often, the target recoils, to a greater or lesser extent, and sits back. Some throws in what appear to be fairly intense social interactions are not directed at another octopus but into empty space.

Although the most obvious possible social role for throwing is that it functions in aggression directed on other animals, another possibility involves the ethological concept of *displacement* (20). This controversial interpretation of some animal behavior holds that animals with elevated arousal, especially frustration, may engage in demonstrative aggressive acts undirected on other individuals, possibly serving as release of the arousal. Some throws in our 2015 data occurred in the midst of, or soon after, intense social interactions but were not apparently directed on other octopuses, and did not hit others. In one case, for example, a male was rebuffed in a mating attempt, and immediately after the rebuff he threw a shell, changed color, and appeared to accelerate his breathing.

A large number of throws were also made by octopuses while at a particular den, where octopuses may have experienced heightened arousal for several reasons. This den, flanked by several others, was the position on the site at which the most social interaction occurred. The most active male visited all the surrounding likely females at various times, and also spent time sitting in the open near this den. This den was also near a camera, and was the den from which most of the camera-directed throws occurred.

Showing intention in a behavior is difficult in non-human animals (12). A deflationary interpretation of the tendency of throws to hit other individuals might be offered as follows: An octopus in a social setting will often attend to other nearby individuals. If an octopus throws debris in the course of den-cleaning or for other non-social reasons, but does so while attending to another, hits may arise fortuitously because of the thrower’s orientation to the other individual. In relation to our data, this interpretation might explain why hits are associated with uniform coloration, but would not explain the tendency for octopuses to use unusual arm arrangements, and to throw silt frequently and algae hardly ever, in cases when they hit other octopuses (compare Figures 2A and 2B). These and other features of the manner in which hits arise in social settings do suggest that some throws are intended to hit others.

Even if no intention to hit other octopuses lies behind these throws, they do have social effects in interactions between individuals at this site. Octopuses can thus definitely be added to the short list of animals who regularly throw or propel objects, and provisionally added to the shorter list of those who direct their throws on other animals. If they are indeed targeted, these throws are directed at individuals of the same population in social interactions – the least common form of nonhuman throwing.

## MATERIALS AND METHODS

The species of octopus at our field site is *Octopus tetricus* Gould, 1852, a medium-sized benthic octopus common in temperate waters around Australia and New Zealand, seen at unusually high densities at a field site in Jervis Bay, NSW, Australia (15, 13). Data was collected using stationary video cameras (GoPro brand, various models) left on tripods at the site, during a combination of recreational diving (2015) and research diving (2016). Three cameras were usually in operation at different parts of the site, with large but incomplete overlaps in field of view. Due to technical problems or disruptive animal behaviors, sometimes only one or two cameras were operating. Video was analyzed later with scoring of behaviors by PGS and DLS, in consultation with the other authors.

A conservative approach was taken to the reidentification of individuals. A few individuals had markings that enable reidentification across breaks in onscreen continuity (SI). General similarity in appearance and behavior was not taken to be sufficient for re-identification.

Sexes were assigned behaviorally, as strict anatomical identification would require considerable interference with octopus behavior. In some cases enlarged suckers were used as an anatomical indicator of male sex. Most males and females were identified by mating behaviors: males made mating attempts by extending arm R3; females accepted, or did not immediately rebuff, these mating attempts. In some cases, an individual was categorized as behaviorally male or female based not on mating behaviors per se, but on other behaviors seen to be associated with male or female mating roles. In the case of males, these behaviors were: interfering with or blocking departure attempts by behavioral females; attending to but not opposing returns from off-site by behavioral females; aggression towards other likely males. In the case of females, these behaviors were avoiding aggression with behavioral males; being interfered with or blocked on departure attempts; being attended to and allowed to approach on return from offsite by behavioral males. A total of 89 of 102 throws were by individuals whose sex could be assigned through observation of mating attempts, based on nominal (more conservative) identifications of individuals.

Throws were scored as high, medium, or low vigor. High vigor throws propelled a mass of material comparable in size to the animal’s body, and/or propelled it a distance comparable to the animal’s body. Silt throws produced larger plumes than other thrown materials; we made allowance for this fact informally. Low vigor throws propelled a much smaller quantity of material – one or a few shells, for example – a distance that was a small fraction of the animal’s body size. Medium vigor throws were intermediate between these. Scoring was not based on quantitative criteria. Borderline cases described in the Results and SI were included in low vigor throws.

One interest of our analyses is to examine to what extent throwing a projectile was an agonistic social tool use, and we thus identified social, mixed and two categories of non-social contexts (den throws and eating, see Results). We structured our analyses to use den and eating throws as field controls and examined whether “control” (non-social) throws differed from social throws.

We took throws to be the behavior of interest, and examined how aspects of throwing change among variable short-term ecological contexts. We used chi-square (for context and body pattern during throws) and Fisher’s exact tests for association (for throw outcomes). Throwers varied in both contexts, patterns, materials thrown and so forth; and context, for example, was a function of the behavior of all octopuses on the site.

## ETHICS STATEMENT

This study comprised unobtrusive observation only of undisturbed wild non-protected invertebrate animals that were not manipulated in any way. The 2016 data collection was approved by the Alaska Pacific University Institutional Review Board and the University of Sydney Animal Ethics Committee.

## AUTHOR CONTRIBUTIONS

PGS, DS, and ML conceived and designed the project. All authors contributed to data collection. PGS and DS analyzed the data, in consultation with SL and SC. PGS and DS wrote the manuscript text. All authors reviewed the manuscript in draft and made comments.

## COMPETING INTERESTS

The authors declare no competing interests.

## ACKNOWLEDGMENTS

We are grateful to the Alaska Pacific University Scientific Dive Program for support. Sue Newson provided logistical support. Research diving in 2016 was conducted under permit BDR16/00008 to PGS, DS, and SC to conduct research in Booderee National Park. Figure 1 panels c and d were prepared by Rebecca Gelernter.

## FINANCIAL STATEMENT

Financial support for this study was provided to PGS by the City University of New York and to DS through Alaska Pacific University from donations by the Pollock Conservation Consortium. Findings and conclusions presented by the authors are their own and do not necessarily reflect the views or positions of the supporting organizations.

## Notes

### Competing Interest Statement

The authors have declared no competing interest.

