## Supporting Information for "In the Line of Fire: Debris Throwing by Wild Octopuses"

Peter Godfrey-Smith, David Scheel, Stephanie Chancellor, Stefan Linquist, Matthew Lawrence

**I: Videos [* Not included in preprint version]**

All videos have had their colors adjusted to increase contrast and the visibility of behaviors.

Video 1: A throw by a female octopus (T1F) that hits a male attempting to mate with her. The material thrown is silt, vigor is high, and thrower's pattern is dark uniform. (2015 data, throw 27-14)

Video 2: A throw by a female octopus (T23F) that hits another likely female. This throw is part of the concentrated sequence of throws with a high rate of hits recorded at the end of the full day of sampling in the 2015 data. The throw is directed from underneath arm L1, rather than between L1 and R1, so it is one of the "anomalous arm" throws discussed in the main text. The material thrown is a combination of shells and silt, vigor is medium, and the thrower's pattern is mottled. (2015 data, throw 28-46)

Video 3: A throw by a female octopus, hitting a behavioral male. The material thrown is silt. The male ducks and raises an arm just before the throw is made. (2016 data, throw 17-11)

Video 4: A throw by a female octopus disposing of fresh shells after eating. The octopus returned from a foraging trip 18 mins earlier and assumed a characteristic feeding posture. As the motion of the throw begins, another octopus reaches towards her and they touch as the shells are released. The context of this throw was scored as Eating (E) despite the reach. The vigor was scored as high and the pattern as mottled. (2015 data, throw 27-19)

Video 5: A throw by a female octopus engaged in den maintainence. In the 2 minutes prior to the throw, the octopus several times was engaged in bringing up shells from inside her den. The throw is directed from between R1 and R2, so it is another "anomalous arm" throw. The throw was scored as medium vigor with a mottled pattern. (2015 data, throw 29-07)

### II: Methods and Further Details of Results

#### 1. Categorization of behaviors

As noted in the main text, throws of the kind described in this paper shade into other behaviors, because the gathering and holding of material can be minimal. Some apparent throws are also made from positions partially inside a den, making observation of behavioral details difficult. We opted to include, rather than ignore, some borderline cases that only minimally met our definition, as very little material was projected, the force of projection was minimal, and/or gathering prior to the throw was minimal. There were 11 (approximately 11% of N=102) such borderline throws; thus they comprise a small portion of our sample.

**2. Time of day at which throws occurred**

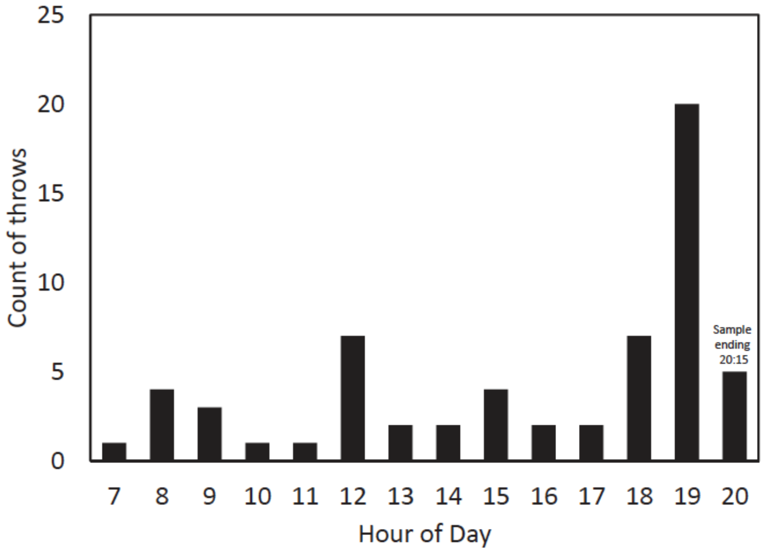


Figure S1: Throws by hour of occurrence during the day. On the day with 13.5 hours of sampling, throws occurred throughout daylight hours but increased in frequency around dusk

**3. Individual identification**

A few individuals had markings that enable reidentification across breaks in onscreen continuity; these were the two most frequent female throwers – T1F, T23F – and, for part of the data collection period, T6M (see Figure S2). It was not possible to track all individuals over long periods, so we did not attempt to determine whether the elevated observed rates of throwing by some individuals may be due to them spending more time on screen. As discussed in the main text, a particular den was very often the site of throws, a finding not affected by difficulties in tracking individuals.

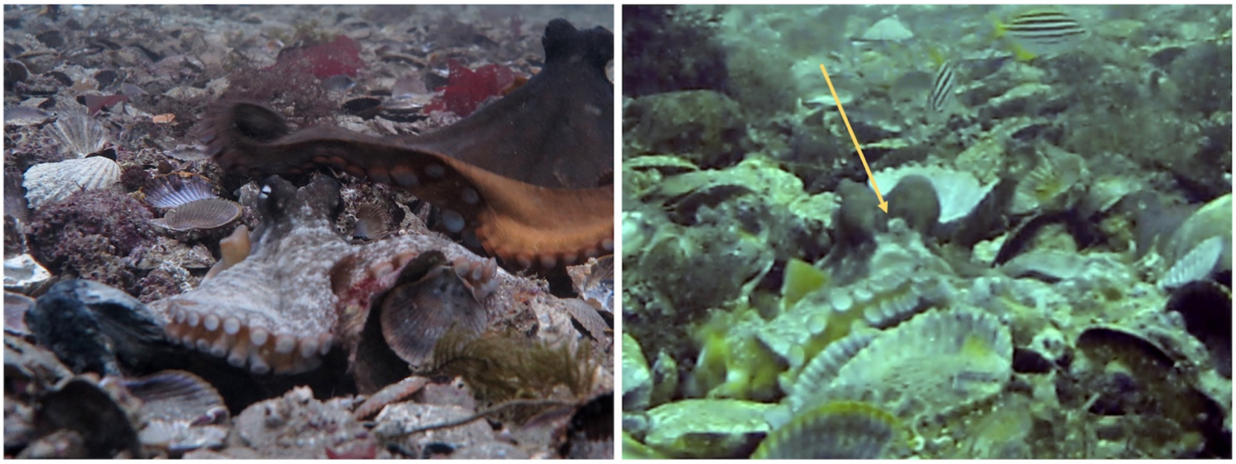

Figure S2: Individually recognizable markings below the eyes of two individuals, TIF (left) and T23F (right). For T1F, note the pale dot (papillae) below each eye higher than the frontal white spot and the prominent and almost unbroken horizontal white frontal bar below the frontal white spot. For T23F, note the arc-shaped or peanut-shaped white area comprising the upper part of the frontal white spot (indicated by the arrow). When the relevant body patterns were displayed by an octopus, these distinctive details were consistently noted on T1F and T23F respectively and were absent on other octopuses. GoPro camera image on the right has been adjusted for color tone. Left image: photo by Peter Godfrey-Smith. Right image: from video data used in this paper.

**4. Contexts of behaviors**

Some throws were difficult to score for context and handled in a way that included consideration of factors outside the 2 minute window, or set aside events within it. In one case, a lengthy quiet mating included a clear eating throw by the female, and we scored this not as mixed, but as an eating throw. In a second case, no indicators of context occurred until after the throw was in process, 5 seconds before release, at which moment another octopus arrived, probably not influencing the throw; this we scored as no-context. In a third case, no relevant behaviors occurred within the 2 minute window but events outside it suggested a den-cleaning context. In a fourth case (see video 4 of SI), during the initial motions of a throw disposing of fresh prey remains after foraging, another octopus reached towards the thrower; this was categorized as an eating throw. Two cases were also ambiguous because, as discussed in the main text, an octopus may have been responding in part to the presence of a fixed camera on a tripod within a meter of its den. Based on other behaviors, both were scored as den-cleaning throws. A total of six cases of 101 were thus ambiguous in these ways.

**5. Details of hits**

In 17 cases, as noted in the text, material thrown hit another octopus. In one additional case, a "hit" resulted as an octopus moved into the cloud after it was thrown. In two additional cases, material thrown hit a fish (minimally in one case). A large majority of the hits on other octopuses were due to the two throwers who threw most frequently in general: 5 hits from T1F and 9 from T23F). The other hits were single cases from three different individuals (probably one male and two female). A total of 15 of 17 hits were by behavioral females as identified by mating behavior, with one additional thrower a likely female due to other behaviors. The sex of the octopus hit by a throw could be assigned behaviorally in 13 cases out of 17. These comprised 8 hits on behavioral females (all sexes assigned by behaviors other than mating) and 5 on males (all assigned by mating, and probably all the most active male discussed in the main text).
